# Chaotropic Agent-assisted Supported Lipid Bilayer Formation

**DOI:** 10.1101/2024.07.02.601713

**Authors:** Jennie L. Cawley, Dane E. Santa, Aarshi N. Singh, Adeyemi T. Odudimu, Brett A. Berger, Nathan J. Wittenberg

## Abstract

Supported lipid bilayers (SLBs) are useful structures for mimicking cellular membranes, and they can be integrated with a variety of sensors. While there are a variety of methods for forming SLBs, many of these methods come with limitations in terms of the lipid compositions that can be employed and the substrates upon which the SLBs can be deposited. Here we demonstrate the use of an all-aqueous chaotropic agent exchange process that can be used to form SLBs on two different substrate materials: SiO_2_, which is compatible with traditional SLB formation by vesicle fusion and Al_2_O_3_, which is not compatible with vesicle fusion. When examined with quartz crystal microbalance with dissipation monitoring, the SLBs generated by chaotropic agent exchange (CASLBs) have similar frequency and dissipation shifts to SLBs formed by the vesicle fusion technique. The CASLBs block nonspecific protein adsorption on the substrate and can be used to sense protein-lipid interactions. Fluorescence microscopy was used to examine the CASLBs, and we observed long-range lateral diffusion of fluorescent probes, which confirmed the CASLBs were composed of a continuous, planar lipid bilayer. Our CASLB method provides another option for forming planar lipid bilayers on a variety of surfaces, including those that are not amenable to the widely used vesicle fusion method.

## INTRODUCTION

Planar phospholipid bilayer membranes formed on solid substrate supports, commonly known as supported lipid bilayers (SLBs), are employed for a wide variety of applications.^1^ Because SLBs mimic many of the essential properties of cellular membranes, they can be used where biophysical properties of the membrane are of interest.^2–5^ Additionally, integration of SLBs with surface-based sensors can be relatively straightforward, depending on the lipid composition and surface characteristics. SLBs can be incorporated into a broad range of sensor types, including optical^6–8^ and electrical^9–11^ sensors, and chemical imaging techniques^12–14^ can be used to investigate SLBs. Therefore, they have found utility as receptor layers for sensors that probe peptide-lipid,^15^ lipid-lipid,^16^ nanoparticle-lipid,^17^ ion-lipid,^18^ and small molecule-lipid^19^ interactions.

The formation of SLBs on solid substrates can be accomplished by a number of different methods that vary in their technical and practical requirements. Sequential deposition of the two bilayer leaflets can be accomplished by Langmuir-Blodgett (L-B) and Langmuir-Shaefer (L-S) methods.^20^ Because the substrate-contacting and distal membrane leaflets are deposited in separate steps, these approaches can be used to create bilayers where the leaflet lipid compositions differ, which facilitates the generation of asymmetric lipid bilayers.^21^ Additionally, L-B and L-S methods can be used to form SLBs with lipid compositions that are difficult to planarize by other methods. An example of this are membranes composed of significant fractions of saturated lipids and cholesterol, including membrane compositions that display coexisting domains with distinct lipid compositions.^22^ Despite the advantages of the L-B and L-S methods, they have some technical limitations, including the necessity of a Langmuir trough and substrate dipping apparatus, which may not be available to all groups.

A more convenient method of SLB formation is the vesicle fusion method. The first step of this approach is to generate phospholipid vesicles. Then, these vesicles are allowed to adsorb onto a substrate. Once a critical surface coverage of vesicles is reached, a vesicle fusion cascade commences, resulting in the formation of a planar SLB.^23^ The rupture process is driven by adhesion between the vesicle membrane and the substrate, along with vesicle-vesicle interactions, and it is opposed by membrane cohesive forces.^24^ The simplicity and convenience of the vesicle fusion method is offset by some limitations. For example, not all substrate materials and lipid compositions are compatible with SLB formation by vesicle fusion.

While vesicles composed primarily of zwitterionic phospholipids, such as phosphatidylcholines (PC), will readily rupture on glass, SiO_2_, and mica surfaces under standard physiological buffer conditions, they will not spontaneously rupture on other oxide surfaces, such as Al_2_O_3_ ^25^ or TiO_2_,^26^ nor will they rupture on bare metallic surfaces such as gold^27^ or silver.^28^ Due to the challenges of SLB formation on many substrate materials, clever SLB formation methods have been developed, including bubble collapse deposition,^29^ peptide-induced vesicle fusion,^30^ tethered lipid bilayers,^31^ and hybrid bilayer formation.^32^ When using substrates amenable to zwitterionic SLB formation, the incorporation of negatively charged lipids, such as gangliosides, can inhibit vesicle fusion.^33^ In addition to negatively charged lipids, lipids with negative spontaneous curvature, such as phosphatidylethanolamine, hinder SLB formation by vesicle fusion,^34^ as do high concentrations of sterols like cholesterol.^35^

The limitations in substrate and lipid composition scope associated with the vesicle fusion method have necessitated the development of other methods to form SLBs. A particularly attractive method of SLB formation suitable with an expanded range of substrates and lipid compositions is the solvent-assisted lipid bilayer (SALB) formation method.^36^ The SALB method builds upon earlier work with lipid membrane assembly during gradual solvent exchange.^37^ It begins with lipids dissolved in an organic solvent like ethanol or isopropanol. The lipid solution in the organic solvent is typically injected into a flow cell attached to a suitable substrate. Then, an aqueous buffer solution is injected into the flow cell to exchange the solvents. After the aqueous phase is injected, the lipids rapidly assemble into a bilayer membrane that deposits on the substrate along the moving organic-aqueous interface.^38^ One major advantage of the SALB method is that it is compatible with a wider range of lipid compositions than vesicle fusion, including compositions possessing extremely high mole fractions of cholesterol.^39^ Moreover, the SALB method can be used to form SLBs on substrates that are not compatible with vesicle fusion, including a variety of oxides and metallic substrates such gold,^40^ which is especially advantageous for plasmonic or electrochemical sensing applications.^41^ While the SALB method has significant advantages, the requirement of organic solvents to solubilize the lipids may limit its applicability in some circumstances.

In an effort to extend the solvent exchange methodology to an all-aqueous system, here we introduce a chaotropic agent-assisted supported lipid bilayer (CASLB) formation method. Chaotropic agents are solutes that cause disorder in biological macromolecules and their assemblies, such as lipid bilayer membranes.^42–43^ Examples of chaotropic agents include small molecules like ethanol, n-butanol, and urea, as well as salts such as guanidinium chloride, sodium tribromoacetate (NaTBA), and sodium trichloroacetate (NaTCA).^44^ These agents exert their effects by a variety of specific mechanisms, but they generally disrupt water-water and water-solute interactions, and thus they have a profound influence on the solubility and structure of a variety of solutes. For example, chaotropic agents can be used to dissociate enzyme complexes^45^ and are used in the purification of hydrophobic membrane proteins, such as caveolin.^46^ Most important for the present studies, chaotropic agents alter phospholipid self-assembly in aqueous solutions. Previous work examining the influence of NaTCA and NaTBA on PC dispersions found that when these salts were present in concentrations greater than approximately 2 M, the turbidity of solution dropped significantly.^47^ This indicates that instead of forming larger assemblies such as vesicles, the PC was primarily in the form of micelles or fully solvated molecules. The low turbidity of PC dispersions was even observed when up to 30 mole % cholesterol was included as part of the lipid mixture. Significantly, when PC is dispersed in an aqueous NaTCA solution and then the solvent is gradually exchanged for chaotrope-free aqueous solution by dialysis, giant vesicles will form.^48^

We wondered if rapid exchange of aqueous solutions of NaTCA for chaotropic agent-free buffer would result in the formation of SLBs on substrates, including substrates that are not compatible with vesicle fusion, such as Al_2_O_3_. We started with a dispersion of PC in 3 M NaTCA, then rapidly exchanged the aqueous medium for a buffer solution inside a flow cell. This resulted in the formation of a SLB on both SiO_2_ and Al_2_O_3_ substrates. We term these SLBs chaotropic agent-assisted SLBs (CASLBs). Herein we describe the lipid concentration requirements for CASLB formation, show that the CASLBs block protein adsorption on the underlying substrate, and demonstrate that CASLBs can be used for detecting protein-lipid binding. Finally, we show that the lipids in the CASLB possess the lateral fluidity that is a hallmark of SLBs.

## MATERIALS AND METHODS

### Chemicals

1-palmitoyl-2-oleoyl-glycero-3-phosphocholine (POPC), 1,2-dipalmitoyl-sn-glycero-3- phosphoenthanolamine-N-(cap biotinyl) sodium salt (biotin-PE) were purchased from Avanti Polar Lipids. Texas Red-dihexadecanoylphosphatiylethanolamine (TR-DHPE), neutravidin, sodium acetate, and buffer salts were purchased from Thermo Fisher Scientific. Sodium tricholoacetate (NaTCA, 97%) was purchased from Alfa Aesar. Bovine serum albumin (heat shock fraction, fatty acid-free) was purchased from Sigma Aldrich. 200 proof ethanol was purchased from Decon Laboratories Inc. All aqueous solutions were prepared with ultrapure water (18.2 MΩ) from a MilliQ system.

### Vesicle and NaTCA lipid solution preparation

Depending on the experiment and the desired lipid composition, lipids (POPC, biotin-cap-PE, TR-DHPE) were dissolved in chloroform and placed in glass vials at desired molar ratios with a total lipid concentration of 1.0 mg/mL. When present in a sample, the biotin-PE and TR-DHPE were held at 1 mole %. The chloroform was evaporated under vacuum in a desiccator at room temperature for a minimum of 2 h. The dry lipid films were rehydrated with aqueous solutions of 3 M NaTCA, 3 M sodium acetate, or Tris buffer (10 mM Tris, 150 mM NaCl, pH 7.0) and gently vortexed. The vesicles resulting from lipid rehydration in Tris buffer were further subjected to 10 min of bath sonication at room temperature, then they were extruded with 23 passes through 50 nm pore diameter polycarbonate filters using a Mini-Extruder (Avanti Polar Lipids) to form vesicles. After extrusion, the vesicles were diluted to the desired total lipid concentration with Tris buffer.

### Quartz crystal microbalance with dissipation monitoring (QCM-D) measurements

A QCM-D (Q-Sense E1 Explorer, Biolin Scientific) utilizing a 5 MHz AT-cut quartz crystal was used for real-time monitoring of the formation of CASLBs or SLBs by vesicle fusion. Silica (SiO_2_) and alumina (Al_2_O_3_)-coated sensor chips (Nanoscience Instruments) were employed for these studies and were extensively cleaned prior to measurements. The SiO_2_ sensor chips were cleaned with a 10 min UV-ozone treatment (ProCleaner Plus, Bioforce Nanosciences), a subsequent soak in 2% (w/v) sodium dodecyl sulfate (SDS) solution, an extensive rinse with ultrapure water, drying with a stream of nitrogen gas, and finally an additional 10 min UV-ozone treatment. The Al_2_O_3_ sensor chips underwent a 10 min UV-Ozone treatment, a 10 min soak in ethanol, an extensive rinse with ultrapure water, drying with a stream of nitrogen gas, and then another 10 min UV-ozone treatment. All measurements were performed at a constant temperature of 23.0 °C and a flow rate of 50 μL/min controlled with a peristaltic pump. Frequency and dissipation signals were monitored at the 3rd, 5th, 7th, 9th and 11th overtones, with analysis performed using the 3rd overtone. To assess if SLBs via CASLB or vesicle fusion were continuous and defect-free, 10 μM BSA in 1× PBS was injected following their formation.

### Neutravidin binding to CASLBs detected with QCM-D

Biotinylated membranes for sensing protein-lipid interactions were composed of POPC and biotin-PE (99:1 molar ratio) and were formed using the CASLB method on SiO_2_ and Al_2_O_3_- coated QCM-D sensor chips. For CASLBs formed on SiO_2_-coated chips, 0.375 mg/mL lipid in 3 M NaTCA was injected, whereas for CASLBs formed on Al_2_O_3_, 0.75 mg/mL lipid in NaTCA was injected. The lipid/NaTCA solutions were infused with a flow rate of 50 μL/min, unless otherwise stated. After formation, CASLBs were washed with Tris buffer. Prior to neutravidin infusion into the QCM-D, the flow rate was increased to 100 μL/min, and 0.1 mg/mL neutravidin was injected. Neutravidin was infused for 20 min and subsequently rinsed with Tris buffer.

### Fluorescence recovery after photobleaching (FRAP)

Bare glass coverslips or coverslips coated with Al_2_O_3_ were used as substrates in the FRAP studies. The Al_2_O_3_ (50 nm thick) was deposited on glass coverslips by atomic layer deposition. Prior to lipid deposition, the bare glass coverslips were cleaned with 2% (w/v) SDS, rinsed with ultrapure water and dried in a stream of nitrogen gas. The Al_2_O_3_-coated coverslips were cleaned by a 10 min exposure to UV-ozone, then soaked in ethanol for 10 minutes, rinsed with ultrapure water and dried in a stream of nitrogen gas. Both SiO_2_ and Al_2_O_3_ coverslips then underwent a 10 min UV-ozone treatment and were bonded to a bottomless 6 channel adhesive chip (sticky-Slide VI 0.4, Ibidi, GmbH). For SLBs formed via vesicle fusion, 0.375 mg/mL POPC liposomes with 1 mole % TR-DHPE in Tris were injected into the channels by hand with a syringe. After 30 minutes, samples were rinsed with Tris buffer. For SLBs formed via CASLB, 0.375 mg/mL POPC lipids with 1 mole % TR-DHPE in 3M NaTCA were injected into the channels via syringe and incubated for 30 mins. The NaTCA solution was exchanged with Tris buffer at a continuous flow rate (50 μL/min) using a syringe pump (Harvard Apparatus 2000). The SLBs formed by vesicle fusion and the CASLBs were examined using an inverted microscope (Nikon Ti) equipped with a 100× 1.49 N.A objective. TR-DHPE fluorescence was excited at 561 nm using an Aura II LED light engine and a standard TRITC filter set. The photobleaching for FRAP was supplied by a 5 s pulse from a 50 mW 405 nm diode laser. Images were captured with a 2048 × 2048-pixel sCMOS camera (Orca Flash 4.0 v2, Hamamatsu). FRAP recovery image stacks were analyzed using the Hankel transform method to calculate the lipid diffusion coefficients.^49^

## RESULTS AND DISCUSSION

### CASLB formation on SiO_2_ and Al_2_O_3_

Our investigation into the CASLB formation process began with a comparison to the vesicle fusion method of SLB formation on SiO_2_ surfaces as well as vesicle adsorption on Al_2_O_3_ surfaces. On SiO_2_ surfaces, the formation of the SLB begins with the adsorption of vesicles on the surface. When observed by QCM-D, the hallmark of vesicle adsorption is a pronounced negative shift in the frequency which is accompanied by an abrupt increase in the dissipation signal. Once a critical population of vesicles has accumulated on the surface, they begin to rupture, at which point the frequency signal is at a minimum and the dissipation is at a maximum. The vesicle fusion cascade continues until eventually a continuous SLB is formed on the surface. At this point the frequency and dissipation signals plateau.^27^ Typical final frequency and dissipation values for a PC SLB on SiO_2_ are between −24 and −26 Hz, and less than 1 × 10^-6^, respectively.^50^ An example of a typical QCM-D trace for vesicle fusion is shown in **Fig. 1a**.

**Figure 1.**
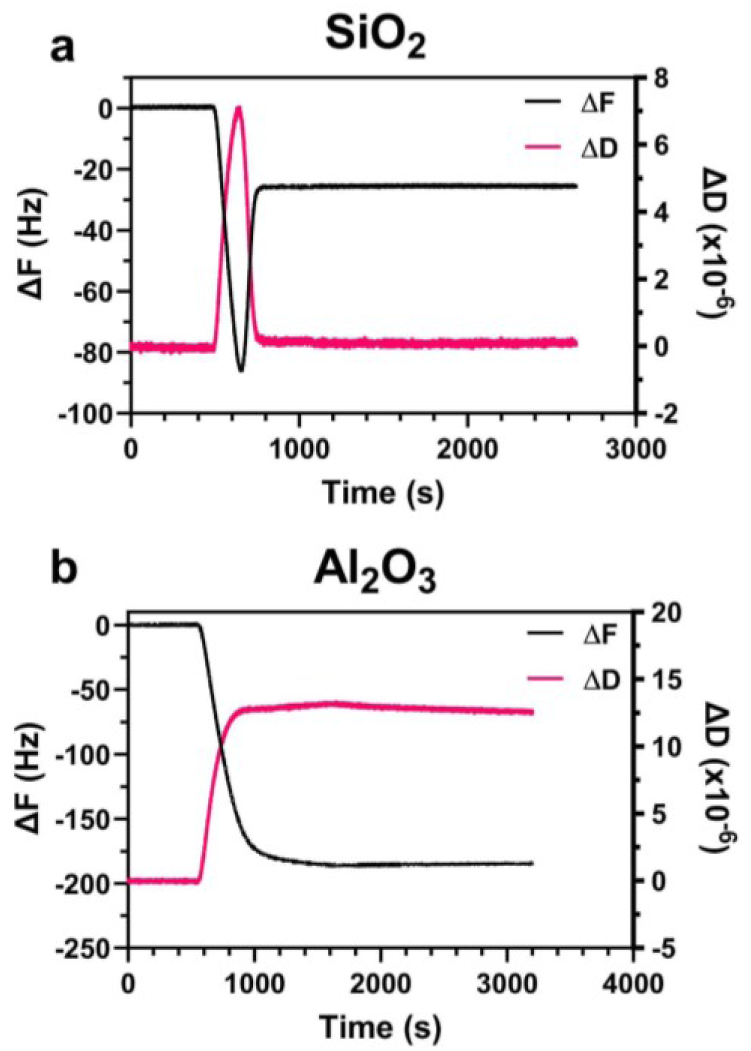
QCM-D response showing the frequency (ΔF, black) and dissipation (ΔD, magenta) shifts for (a) POPC vesicle fusion to form a SLB on SiO_2_ and (b) adsorption of intact POPC vesicles on Al_2_O_3_ substrates.

While vesicles readily adsorb, rupture, and form a SLB on SiO_2_, they adsorb and remain intact on Al_2_O_3_.^29^ In the QCM-D trace, intact vesicle adsorption on Al_2_O_3_ appears as a significant negative shift in the frequency that plateaus at roughly −190 Hz, which is much more negative than the final frequency shift observed for SLB formation on SiO_2_. Coupled with the large negative frequency shift is a large increase in dissipation that plateaus at roughly 13 × 10^-6^. The combination of large negative frequency shift with the large dissipation shift, shown in **Fig. 1b**, is indicative of intact vesicle adsorption, which agrees with previous reports.

For the CASLB formation process, we first dispersed POPC in 3 M NaTCA. In this solution the POPC formed particles with an average diameter of 12.2 nm measured by dynamic light scattering, which is much smaller than vesicles formed upon dispersion of POPC in physiological buffers. After characterizing the POPC dispersions in 3 M NaTCA, we examined the CASLB formation process with QCM-D. Compared to the vesicle fusion method, the CASLB method displays strikingly different frequency and dissipation profiles, owing to the sequential injections of PBS buffer, 3 M NaTCA, 3 M NaTCA with POPC, and finally Tris buffer. An illustration of the steps in the CASLB formation process is shown in **Fig. 2**.

**Figure 2.**
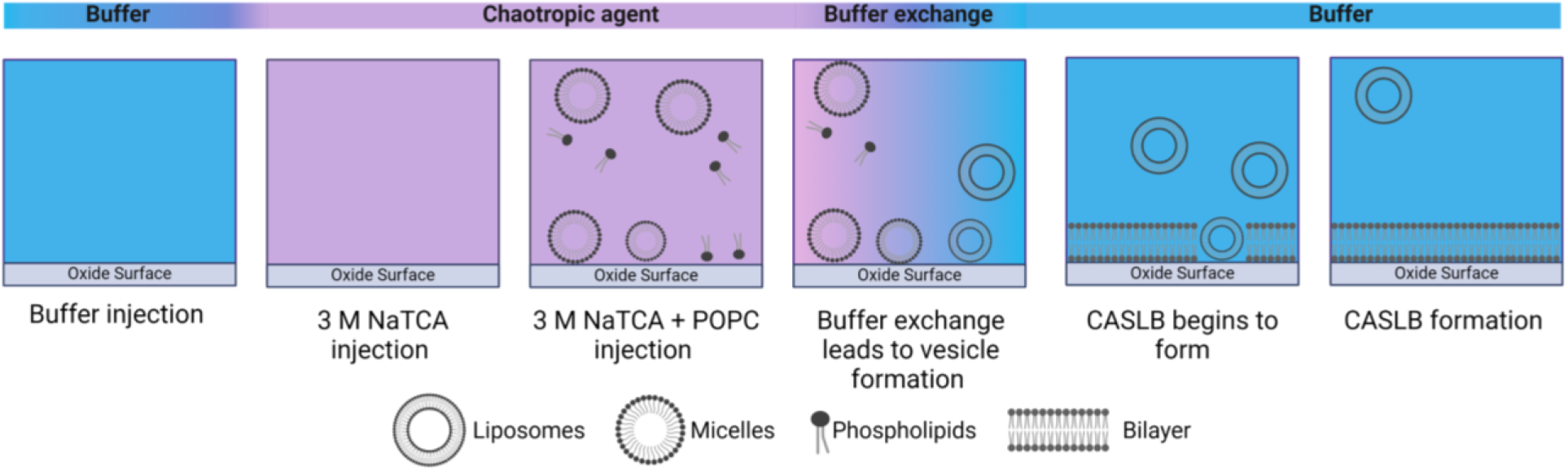
Schematic illustration of the steps in the CASLB formation process.

Annotated QCM-D traces for CASLB formation on SiO_2_ and Al_2_O_3_ are shown in **Fig. 3a**. The annotations in **Fig. 3a** indicate the time points at which the following steps occurred: 1) infusion of PBS buffer and recording of a stable baseline; 2) infusion of 3 M NaTCA; 3) infusion of 3 M NaTCA containing 0.375 mg/mL POPC (SiO_2_) or 0.750 mg/mL (Al_2_O_3_); 4) infusion of Tris buffer; 5) recording of stable frequency and dissipation. All infusions had flow rates of 50 µL/min. The abrupt changes in frequency and dissipation upon infusion of 3 M NaTCA and Tris buffer (steps 2 and 4) are due to the large density differences between 3 M NaTCA and the buffers injected before and after the 3 M NaTCA. We measured 3 M NaTCA to have a density of 1.298 ± 0.043 g/mL compared to 1.001 ± 0.006 g/mL and 1.012 ± 0.011 g/mL for PBS and Tris, respectively. Because they are due to solution density changes, these massive shifts in frequency and dissipation at steps 2 and 4 were observed even when POPC was not included in the 3 M NaTCA solution (dashed lines in **Fig. 3**). However, when POPC was present in the 3 M NaTCA solution, upon the final infusion of Tris, the frequency and dissipation signals did not return to the initial baseline. Instead, residual frequency and dissipation shifts were observed with both SiO_2_ and Al_2_O_3_ substrates. (**Fig. 3b,d**). The residual frequency and dissipation shifts observed after exchanging the NaTCA/POPC solution with Tris buffer indicate that a lipid layer has deposited on the substrate, and we posit that these layers are CASLBs.

**Figure 3.**
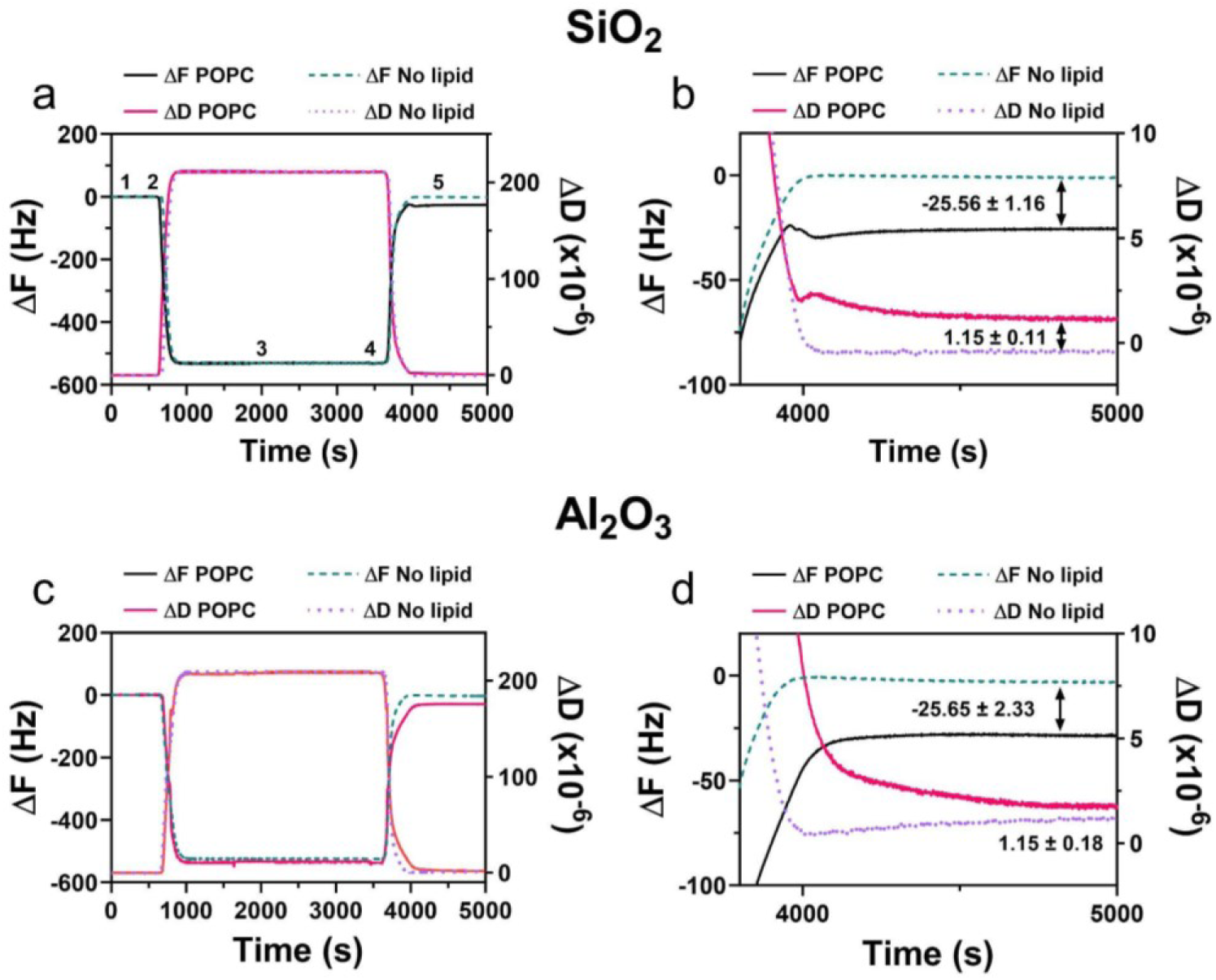
QCM-D frequency (ΔF) and dissipation (ΔD) traces for CASLB formation on SiO_2_ (a-b) and Al_2_O_3_ (c-d) substrates. The full time course of the CASLB formation process is shown in (a) and (c). Steps 1-5 in (a) correspond to 1) PBS buffer infusion; 2) 3 M NaTCA infusion; 3) Infusion of 3 M NaTCA with 0.375 mg/mL POPC; 4) infusion of Tris buffer; 5) Stable signal associated with the formation of a CASLB. (b,d) Magnified view of ΔF and ΔD signals during the final NaTCA-Tris buffer exchange process.

The final frequency and dissipation shifts associated with the deposition of the lipid layer are important figures of merit associated with SLB formation on QCM-D sensors. As mentioned above, POPC SLBs formed by vesicle fusion give rise to frequency shifts in the neighborhood of −25 Hz and dissipation shifts that are less than 1 × 10^-6^. The residual frequency and dissipation values after the final Tris buffer wash in the CASLB method are similar to these ideal values. A summary of the final frequency and dissipation values for lipid layers (SLBs and adsorbed vesicles) on SiO_2_ and Al_2_O_3_ surfaces is shown in **Table 1**. The final frequency and dissipation shifts for the CASLB method on both SiO_2_ and Al_2_O_3_ indicate that the lipid layer deposited is indeed a SLB.

**Table 1.** Summary of QCM-D frequency and dissipation shifts.

| Method | Surface | Final $\Delta F$ (Hz) | Final $\Delta D$ ($\times 10^{-6}$) |
| --- | --- | --- | --- |
| Vesicle fusion | SiO<sub>2</sub> | -25.69 ± 0.15 | 0.10 ± 0.02 |
| Vesicle adsorption | Al<sub>2</sub>O<sub>3</sub> | -189.6 ± 4.50 | 12.97 ± 0.17 |
| CASLB | SiO<sub>2</sub> | -25.56 ± 1.16 | 1.15 ± 0.11 |
| CASLB | Al<sub>2</sub>O<sub>3</sub> | -25.65 ± 2.33 | 1.15 ± 0.18 |

### Effect of lipid concentration on CASLB formation

The data in **Fig. 3** and **Table 1** shows that the CASLB formation process generates membranes with acceptable frequency and dissipation shifts when the infused lipid concentration is 0.375 mg/mL (on SiO_2_) and 0.75 mg/mL (on Al_2_O_3_) in 3 M NaTCA. Next, we sought to determine how tolerant the CASLB formation process is to variations in the infused lipid concentration. To do this, we prepared a series of POPC solutions in 3 M NaTCA where the lipid concentration ranged from 0.125 to 1.0 mg/mL (0.164 mM to 1.32 mM). These solutions were introduced into the QCM-D flow cell over SiO_2_ and Al_2_O_3_ surfaces using the same infusion schedule as described above. After the final Tris buffer infusion step, we measured the final frequency and dissipation shifts, which are shown in **Figure 4**.

**Figure 4.**
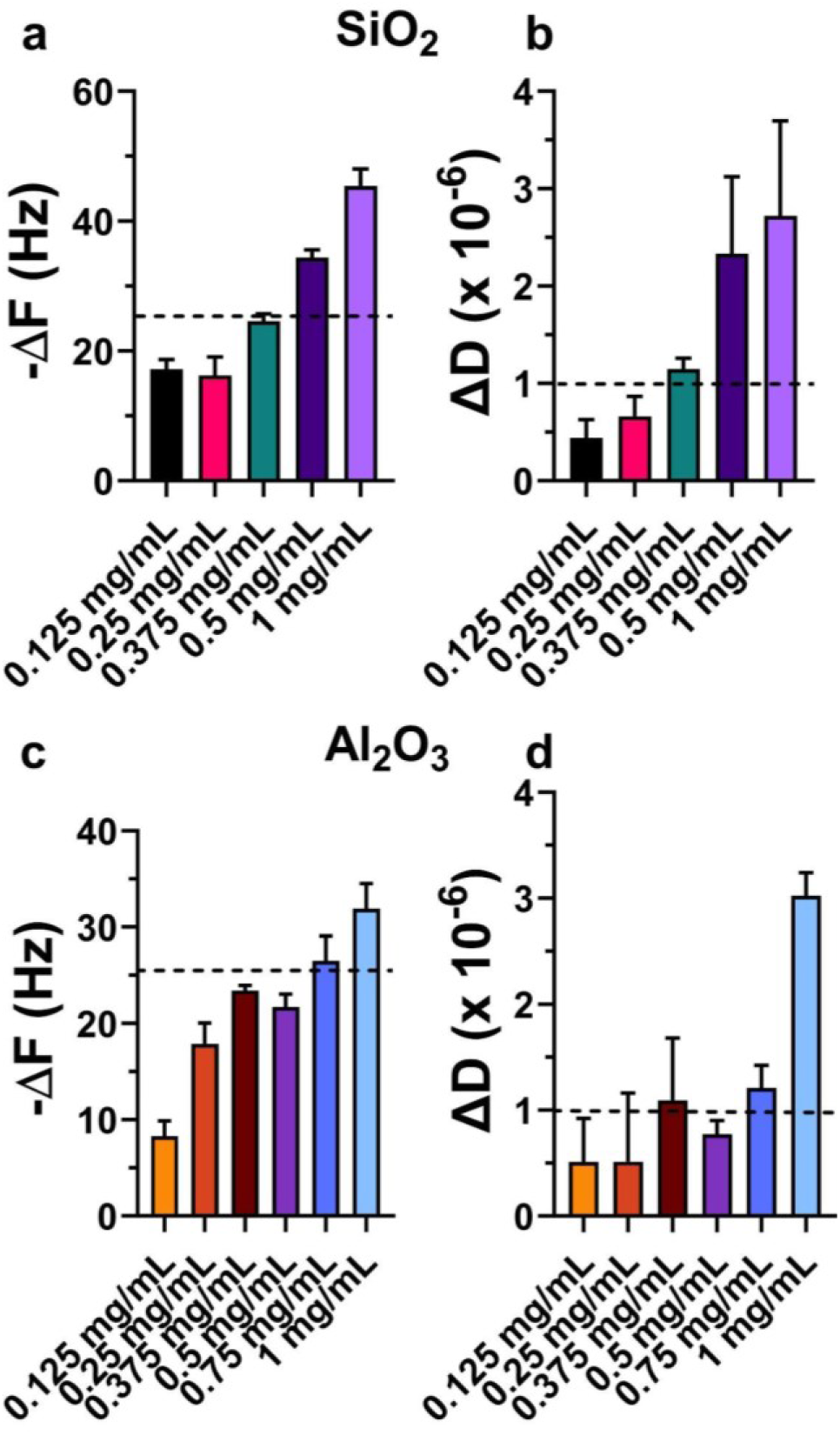
Effect of lipid concentration on the final frequency (ΔF) and dissipation (ΔD) shifts for POPC bilayer formation by the CASLB process on SiO_2_ (a,b) and Al_2_O_3_ (c,d) surfaces. The dashed lines in (a,c) are drawn at −26 Hz and (b,d) 1 × 10^-6^ to indicate typical values for SLBs. Bars heights are mean values for three replicates, and the error bars represent the standard deviations.

Our results indicate that the CASLB formation process is indeed sensitive to the lipid concentration. On SiO_2_ surfaces, 0.375 mg/mL gave frequency and dissipation shifts within the acceptable range, while on Al_2_O_3_ surfaces 0.75 mg/mL was the most suitable. However, 0.375 mg/mL and 0.5 mg/mL also gave acceptable results on Al_2_O_3_. Thus, it appears that CASLB formation on Al_2_O_3_ is somewhat more tolerant to varying lipid concentration. On both surfaces, when suboptimal lipid concentrations were used, the frequency shifts did not attain as large values, indicating that an incomplete bilayer was formed. On the other hand, with greater than optimal lipid concentrations, the frequency shifts were more negative and the dissipations were larger. This could be indicative of multilayers and/or adsorbed vesicular structures on top of the bilayer. Tabaei and coworkers observed similar trends when using the SALB method in organic solvents.^51^

### Trichloroacetate is required for CASLB formation

Next we sought to determine if the trichloroacetate anion of NaTCA was necessary for SLB formation. In this set of experiments, 3 M NaTCA was replaced with 3 M sodium acetate. Previously, it was reported that unlike NaTCA solutions, concentrated sodium acetate solutions do not solubilize PC,^47^ so we suspected that lipid adsorption on SiO_2_ and Al_2_O_3_ surfaces would be altered, and SLBs might not be formed. The lipid concentrations used in these experiments were 0.375 mg/mL for SiO_2_ surfaces and 0.75 mg/mL for Al_2_O_3_ surfaces, and the buffer and lipid infusion schedule was identical to that used for CASLB formation with 3 M NaTCA. Representative QCM-D results are shown in **Fig. 5**. Just like with 3 M NaTCA, the infusion of 3 M sodium acetate resulted in a large shift in the frequency and dissipation signals when using SiO_2_ and Al_2_O_3_ sensors. In a departure from what was observed with NaTCA, with both sensor surfaces a linear decrease in frequency from approximately −125 to −150 Hz was observed while solutions of sodium acetate with POPC flowed over the sensor. Concomitant with this negative frequency shift was an increase in dissipation from roughly 50 × 10^-6^ to 55 × 10^-6^. These two signals are signs of appreciable lipid adsorption on the sensor.

**Figure 5.**
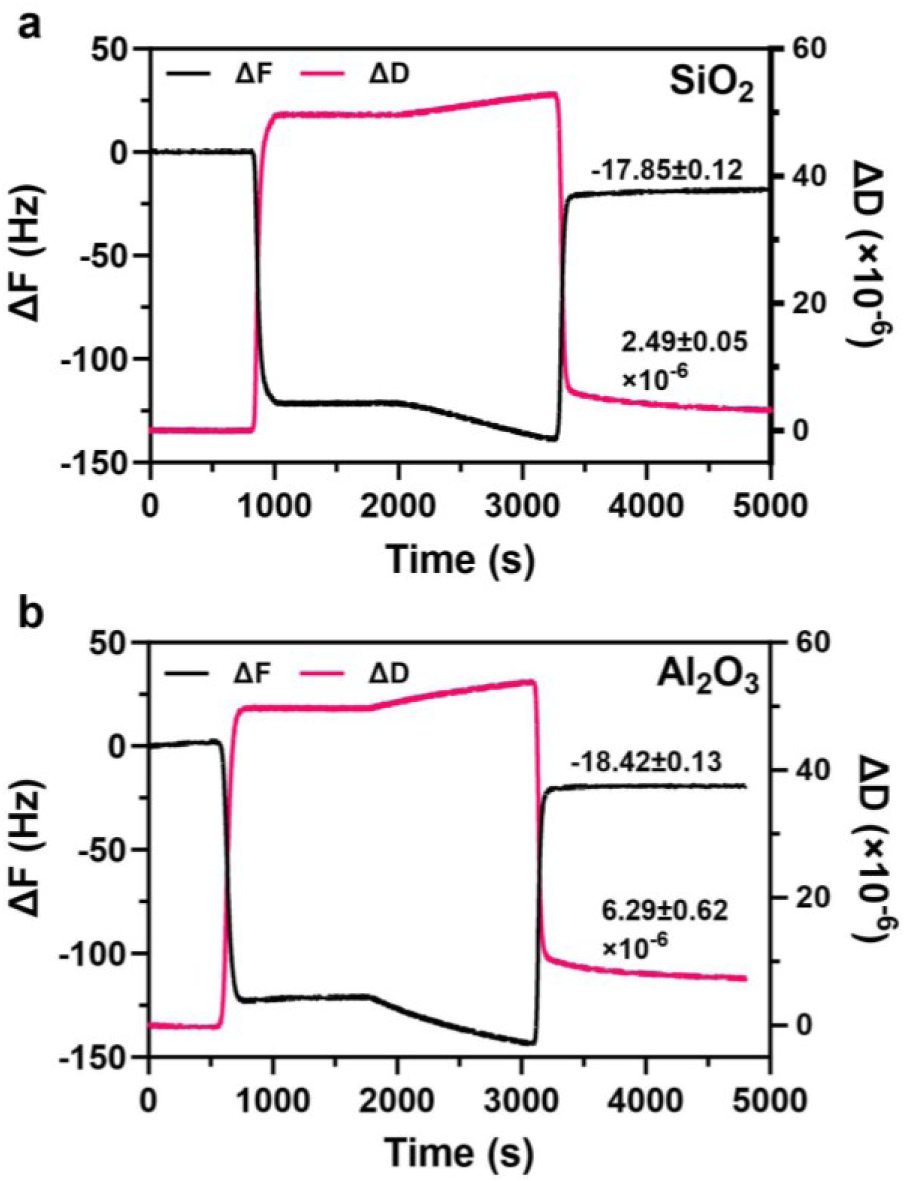
CASLB process requires NaTCA. QCM-D traces showing frequency (ΔF, black) and dissipation (ΔD, magenta) recorded with Al_2_O_3_ (a) and SiO_2_ (b) coated sensors when POPC is infused in 3 M sodium acetate instead of 3 M NaTCA.

Once the Tris buffer was infused in the last step of the process, the frequency stabilized in −17 to −19 Hz range, along with dissipation shifts that plateaued between 2 × 10^-6^ and 7 × 10^-6^. The final frequency shift values, which are less negative than those observed for POPC CASLBs formed using NaTCA, could be indicative of incomplete SLB formation. However, the observation of larger than optimal dissipation shifts suggests the surface layer may be of the form of sparsely adsorbed vesicles. Taken together, these results suggest that the chaotropic anion TCA is a necessary component for successful SLB formation.

### CASLB membranes block nonspecific protein adsorption

A wide variety of biomolecular interactions can be examined using SLBs as receptor layers on QCM-D sensors.^52^ For these studies to be successful, nonspecific interactions should be minimized. Significant nonspecific interactions between the biomolecule of interest and the sensor can occur if the SLB has defects that leave the sensor surface exposed. To determine how well CASLBs block nonspecific adsorption, we used bovine serum albumin (BSA) adsorption to probe the CASLBs for defects. A defect-free SLB will have minimal BSA adsorption. On the other hand, BSA will adsorb to exposed oxide surfaces present within defects. After forming the CASLB on SiO_2_ or Al_2_O_3_ surfaces, BSA (10 µM) was injected into the flow cell and the frequency and dissipation were monitored. **Fig. 6a** shows the frequency shifts as a function of time for BSA adsorption on bare SiO_2_ and bare Al_2_O_3_, as well as on CASLBs formed on SiO_2_ and Al_2_O_3_. From these curves, it is readily apparent that the CASLBs block most BSA adsorption. SLBs on SiO_2_ formed by vesicle fusion also successfully block BSA adsorption **(Fig. S1)**. The average frequency shifts for BSA on all surfaces is summarized in **Fig. 6b**. While the frequency shift for BSA adsorption on CASLBs on SiO_2_ is negligible, there is a small amount of adsorption (ΔF = −3.6 Hz) on the CASLBs on Al_2_O_3_. This suggests that there may be a few small defects in the CASLBs on Al_2_O_3_ substrates.

**Figure 6.**
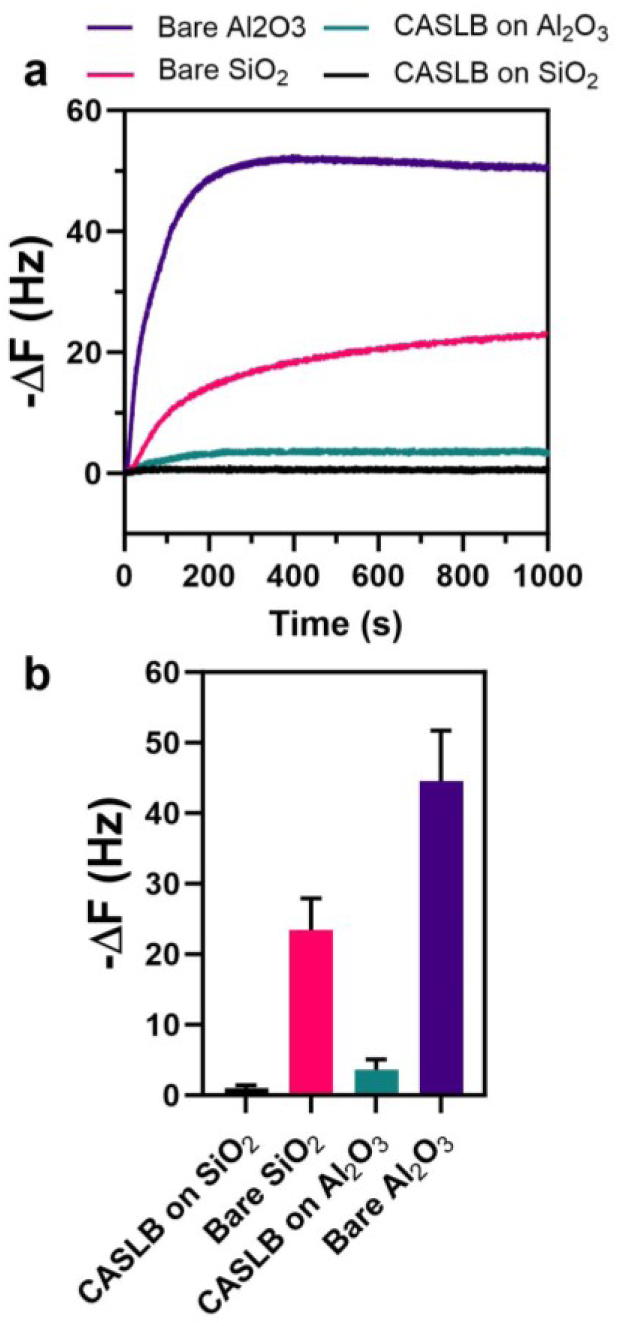
CASLBs inhibit nonspecific protein adsorption. (a) Frequency shift (ΔF) associated with BSA adsorption on POPC CASLB membranes and on bare SiO_2_ and Al_2_O_3_. (b) Average frequency shifts observed for BSA adsorption on the various surfaces. Bars heights are mean values for three replicates, and the error bars represent the standard deviations.

### Detection of protein-lipid binding on CASLB membranes

After determining that the CASLB membranes block nonspecific protein adsorption to sensors, we next sought to use them for monitoring specific protein-lipid interactions. As a model system, we investigated neutravidin binding to biotinylated phosphatidylethanolamine (biotin-PE, 1 mole %) doped into POPC membranes. In these experiments, CASLBs were formed on SiO_2_ and Al_2_O_3_ surfaces, and then neutravidin (0.10 mg/mL) was injected for 20 minutes. As shown in **Fig. 7a**, neutravidin readily binds to the CASLB membranes formed on SiO_2_. When biotin-PE is absent from the CASLB, neutravidin does not bind, as expected. Additionally, there is no appreciable neutravidin dissociation upon injection of pure buffer, due to the extremely strong interaction between biotin and neutravidin. On an Al_2_O_3_ surface, neutravidin also readily binds the CASLB possessing biotin-PE **(Fig. 7b)**. However, there is a non-negligible amount of neutravidin binding to the CASLB that lacks biotin-PE. This is likely related to pinhole defects in the CASLB when prepared on Al_2_O_3_ surfaces. In **Fig. 6** we showed that some nonspecific BSA adsorption occurs when the CASLB is on an Al_2_O_3_ surface. It is likely that similar nonspecific adsorption is occurring when neutravidin is injected over a CASLB on an Al_2_O_3_ surface. These results show that trace lipids can successfully be incorporated in CASLBs and be used to detect protein-lipid interactions.

**Figure 7.**
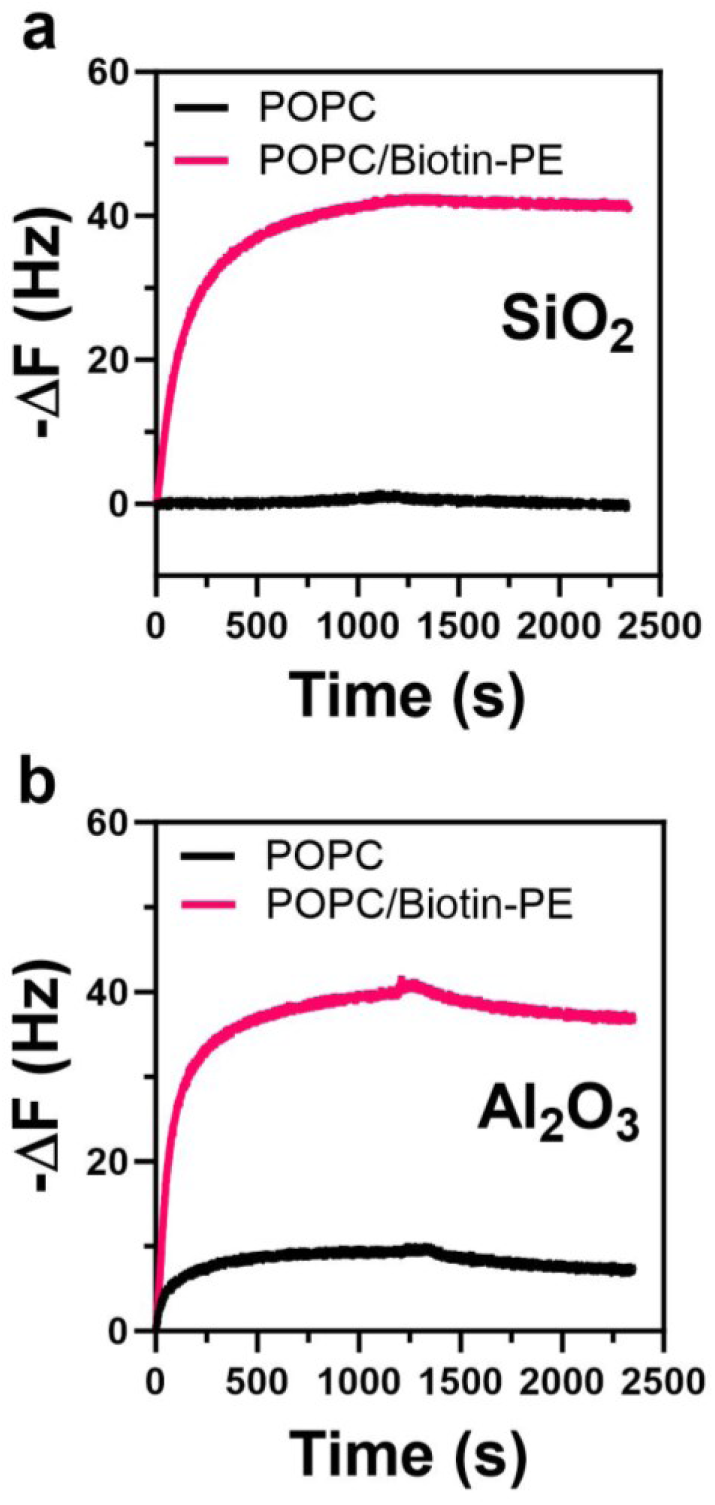
Neutravidin binding to biotinylated CASLB membranes. (a) Frequency shift (ΔF) due to neutravidin binding to a biotinylated CASLB (magenta) or biotin-free CASLB (black) on a SiO_2_ surface. (b) Frequency shift (ΔF) due to neutravidin binding to a biotinylated CASLB (magenta) or biotin-free CASLB (black) on an Al_2_O_3_ surface.

### Lipid diffusion in CASLB membranes

Lateral diffusion of lipids is a hallmark of a continuous planar SLB. Here, we employed fluorescence recovery after photobleaching (FRAP) to examine lipid diffusion in CASLBs compared to SLBs formed by vesicle fusion. In these studies, the lipid composition was POPC with 1 mole % of TR-DHPE as a fluorescent lipid probe. First, lipid vesicles were deposited on bare glass (SiO_2_) and Al_2_O_3_-coated substrates using a microfluidic flow cell. Representative fluorescence micrographs of these lipid layers are shown in **Fig. 8a**. On SiO_2_, the photobleached spots recover within 30 s, indicating that a fluid SLB is formed because the deposited vesicles spontaneously rupture. On the other hand, with Al_2_O_3_ substrates the photobleached spot does not recover. Adsorbed vesicle layers do not display fluorescence recovery after photobleaching because the intact vesicles prohibit long-range diffusion of fluorescent lipid probes. The formation of an adsorbed intact vesicle layer on Al_2_O_3_ agrees with previous reports.^29^

**Figure 8.**
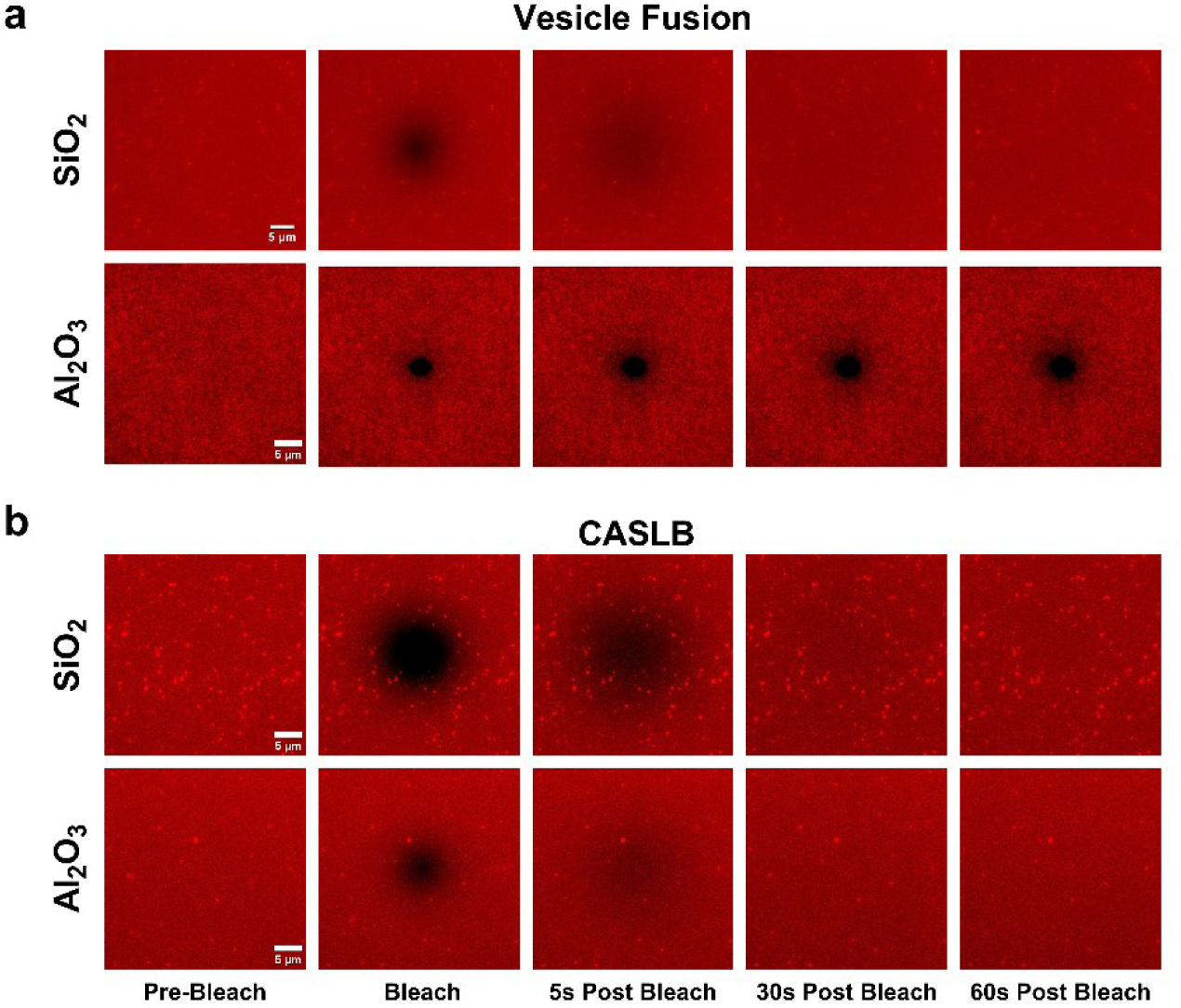
Fluorescence recovery after photobleaching. (a) Fluorescence images of SLBs on SiO_2_ prepared by vesicle fusion (top) and adsorbed intact vesicles on Al_2_O_3_ (bottom). (b) Fluorescence images of CASLBs prepared on SiO_2_ (top) and Al_2_O_3_ (bottom).

Like the SLBs formed by vesicle fusion on SiO_2_ substrates, CASLBs on SiO_2_ displayed fluorescence recovery, indicating that a continuous, fluid membrane was formed **(Fig. 8b)**. On Al_2_O_3_ substrates, FRAP results show striking differences between adsorption of intact vesicles and CASLBs. We observed fluorescence recovery with the CASLBs on Al_2_O_3_, which is strong evidence that a planar supported bilayer is formed by the chaotropic agent exchange process. For CASLBs on both SiO_2_ and Al_2_O_3_ there tended to be a few residual vesicles adsorbed to the membrane after washing. These vesicles may be responsible for the slightly larger than ideal dissipation of CASLBs detected in the QCM-D experiments **(Table 1)**.

Fluorescence recovery curves for the CASLB membranes are shown in **Fig. 9a**. Calculated diffusion coefficients are shown in **Fig. 9b**. The diffusion coefficients for CASLBs were 3.39 ± 0.25 μm^2^/s and 2.19 ± 0.62 μm^2^/s (mean ± s.d.) on SiO_2_ and Al_2_O_3_ substrates, respectively. For comparison, SLBs formed by vesicle fusion on SiO_2_ had a diffusion coefficient of 3.18 ± 0.22, which was not statistically different from the CASLBs on SiO_2_. Because vesicles did not rupture to form a SLB on Al_2_O_3_ surfaces, the diffusion coefficient was not calculated. Our results agree with previous observations of smaller diffusion coefficients in SLBs on Al_2_O_3_ compared to SiO_2_.^53^

**Figure 9.**
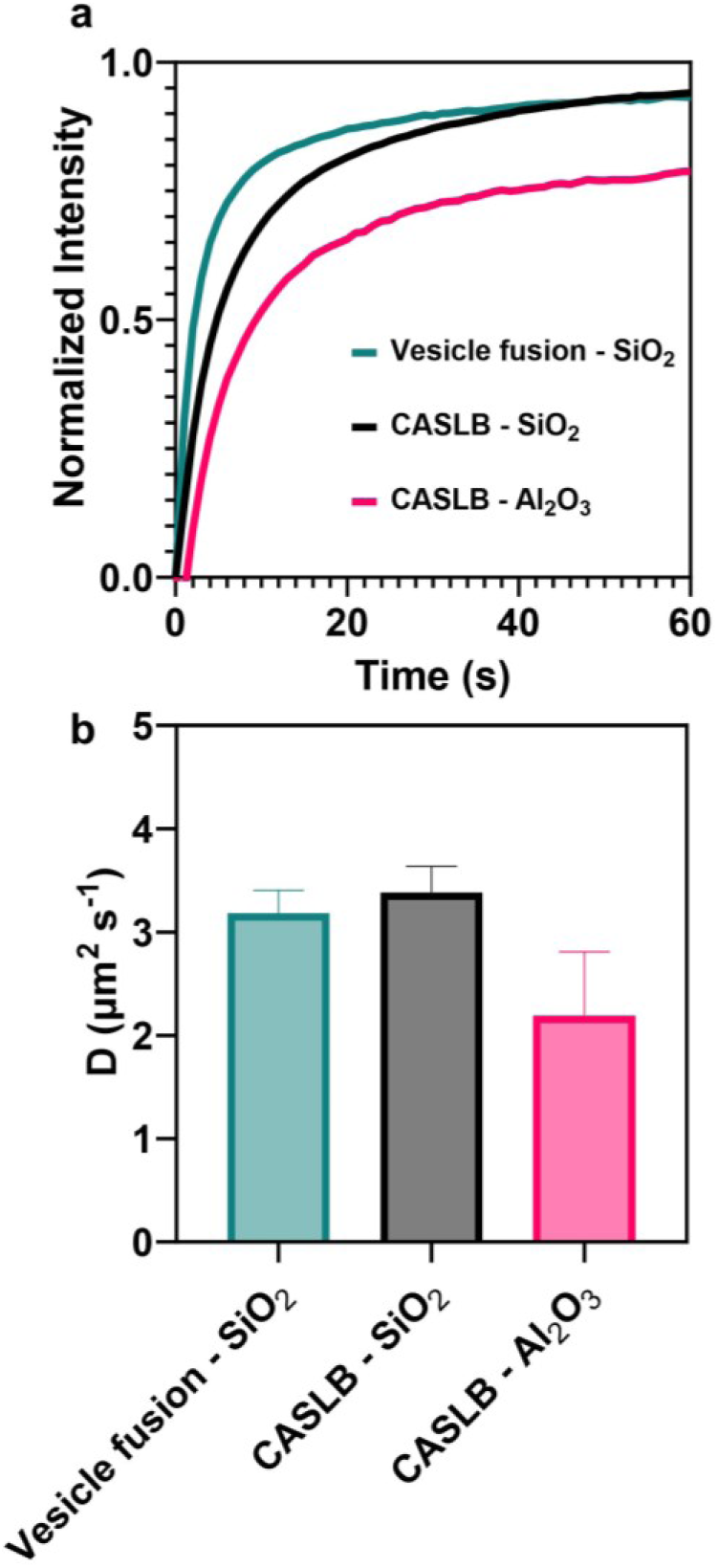
Summary of FRAP results. (a) FRAP recovery curves of SiO_2_ and Al_2_O_3_ slides. (b) Lipid diffusion coefficients in SLBs prepared by vesicle fusion and CASLBs on SiO_2_ and Al_2_O_3_ substrates.

Additionally, the immobile fractions were greater (lower maximum recovery) on Al_2_O_3_ than they were on SiO_2_. A larger immobile fraction could result from a greater number of intact vesicles adsorbed on top of the CASLB on Al_2_O_3_, however, this was not observed. Instead, the smaller diffusion coefficient and larger immobile fraction on Al_2_O_3_ is likely due to a combination of factors, such as hydrodynamic coupling and membrane-substrate interaction energy.^53^ Water has been shown to be more tightly coupled to Al_2_O_3_ than to SiO_2_.^54–55^ The reduced mobility of the water in the hydration layer^56^ between the CASLB and the Al_2_O_3_ substrate could, in effect, more tightly couple the CASLB to the substrate thereby reducing diffusive mobility. Additionally, attractive van der Waals forces between a PC membrane and Al_2_O_3_ are stronger than they are between the same membrane and SiO_2_.^53^ The stronger membrane-substrate interactions should also reduce the diffusive mobility of the lipids comprising the membrane.

## CONCLUSIONS

The formation of SLBs by simple methods, such as vesicle fusion, is somewhat limited in the range of lipid compositions and substrates that can be used. Here we introduce a new way to form SLBs based on the exchange of a chaotropic agent NaTCA. Unlike the solvent-assisted lipid bilayer formation method, our approach does not involve organic solvents and is an entirely aqueous process. We show that similar to vesicle fusion, the CASLB formation method can successfully generate SLBs on SiO_2_ substrates. In contrast to vesicle fusion, however, the CASLB method can also form SLBs on Al_2_O_3_. We show that the CASLBs have QCM-D frequency and dissipation shifts indicative of supported bilayer formation. The CASLBs have few defects and can block nonspecific protein adsorption, and they can also be used for the detection of specific protein-lipid interactions. Furthermore, the CASLB membranes display the lateral fluidity that is a hallmark of planar supported bilayers. While we only investigated oxide surfaces here, we suspect that our approach could be translated to the formation of supported bilayers on different materials, such as metallic or semiconducting substrates. Through the elimination of organic solvents, our approach could potentially be used for a “one-shot” approach for the self-assembly of supported bilayers that contain complex lipid mixtures or incorporate membrane anchored proteins.

## SUPPORTING INFORMATION

Additional QCM-D data (PDF).

## ACKNOWLEDGEMENTS

This work was supported by a grant from the National Science Foundation to N.J.W. (Award number 2044792) and Lehigh University. D.E.S. was supported by a National Science Foundation Graduate Research Fellowship. The authors wish to acknowledge Sang-Hyun Oh and Daniel Upcraft for deposition of Al_2_O_3_ on glass substrates. Illustrations were created with Biorender.

---

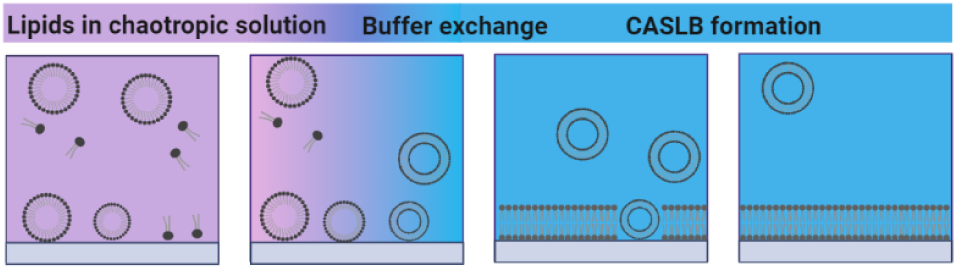
TOC Graphic.

## REFERENCES

1. Castellana, E. T.; Cremer, P. S., Solid supported lipid bilayers: From biophysical studies to sensor design. Surf Sci Rep 2006, 61 (10), 429–444.

2. Parthasarathy, R.; Yu, C. H.; Groves, J. T., Curvature-modulated phase separation in lipid bilayer membranes. Langmuir 2006, 22 (11), 5095–9.

3. Ryu, Y. S.; Yoo, D.; Wittenberg, N. J.; Jordan, L. R.; Lee, S. D.; Parikh, A. N.; Oh, S. H., Lipid Membrane Deformation Accompanied by Disk-to-Ring Shape Transition of Cholesterol-Rich Domains. J Am Chem Soc 2015, 137 (27), 8692–5.

4. Soloviov, D.; Cai, Y. Q.; Bolmatov, D.; Suvorov, A.; Zhernenkov, K.; Zav’yalov, D.; Bosak, A.; Uchiyama, H.; Zhernenkov, M., Functional lipid pairs as building blocks of phase-separated membranes. Proc Natl Acad Sci U S A 2020, 117 (9), 4749–4757.

5. Sapuri-Butti, A. R.; Wang, L.; Tetali, S. D.; Rutledge, J. C.; Parikh, A. N., Interactions of different lipoproteins with supported phospholipid raft membrane (SPRM) patterns to understand similar in-vivo processes. Biochim Biophys Acta Biomembr 2021, 1863 (3), 183535.

6. Jonsson, M. P.; Jonsson, P.; Dahlin, A. B.; Hook, F., Supported lipid bilayer formation and lipid-membrane-mediated biorecognition reactions studied with a new nanoplasmonic sensor template. Nano Lett 2007, 7 (11), 3462–8.

7. Rodrigo, D.; Tittl, A.; Ait-Bouziad, N.; John-Herpin, A.; Limaj, O.; Kelly, C.; Yoo, D.; Wittenberg, N. J.; Oh, S. H.; Lashuel, H. A.; Altug, H., Resolving molecule-specific information in dynamic lipid membrane processes with multi-resonant infrared metasurfaces. Nat Commun 2018, 9 (1), 2160.

8. Yin, H.; Mensch, A. C.; Lochbaum, C. A.; Foreman-Ortiz, I. U.; Caudill, E. R.; Hamers, R. J.; Pedersen, J. A., Influence of Sensor Coating and Topography on Protein and Nanoparticle Interaction with Supported Lipid Bilayers. Langmuir 2021, 37 (7), 2256–2267.

9. Wiegand, G.; Arribas-Layton, N.; Hillebrandt, H.; Sackmann, E.; Wagner, P., Electrical properties of supported lipid bilayer membranes. Journal of Physical Chemistry B 2002, 106 (16), 4245–4254.

10. Zhou, F.; Pan, W.; Chang, Y.; Su, X.; Duan, X.; Xue, Q., A Supported Lipid Bilayer-Based Lab-on-a-Chip Biosensor for the Rapid Electrical Screening of Coronavirus Drugs. ACS Sens 2022, 7 (7), 2084–2092.

11. Duan, Y.; Chen, J.; Jin, Y.; Tu, Q.; Wang, S.; Xiang, J., Antibody-Free Determinations of Low-Mass, Soluble Oligomers of Abeta(42) and Abeta(40) by Planar Bilayer Lipid Membrane-Based Electrochemical Biosensor. Anal Chem 2021, 93 (7), 3611–3617.

12. Pace, H. P.; Sherrod, S. D.; Monson, C. F.; Russell, D. H.; Cremer, P. S., Coupling supported lipid bilayer electrophoresis with matrix-assisted laser desorption/ionization-mass spectrometry imaging. Anal Chem 2013, 85 (12), 6047–52.

13. Pandey, Y.; Kumar, N.; Goubert, G.; Zenobi, R., Nanoscale Chemical Imaging of Supported Lipid Monolayers using Tip-Enhanced Raman Spectroscopy. Angew Chem Int Ed Engl 2021, 60 (35), 19041–19046.

14. Grusky, D. S.; Moss, F. R., 3rd; Boxer, S. G., Recombination between (13)C and (2)H to Form Acetylide ((13)C(2)(2)H(-)) Probes Nanoscale Interactions in Lipid Bilayers via Dynamic Secondary Ion Mass Spectrometry: Cholesterol and GM(1) Clustering. Anal Chem 2022, 94 (27), 9750–9757.

15. Nelson, N.; Opene, B.; Ernst, R. K.; Schwartz, D. K., Antimicrobial peptide activity is anticorrelated with lipid a leaflet affinity. PLoS One 2020, 15 (11), e0242907.

16. Kunze, A.; Bally, M.; Hook, F.; Larson, G., Equilibrium-fluctuation-analysis of single liposome binding events reveals how cholesterol and Ca2+ modulate glycosphingolipid trans-interactions. Sci Rep 2013, 3, 1452.

17. Lee, K.; Zhang, L.; Yi, Y.; Wang, X.; Yu, Y., Rupture of Lipid Membranes Induced by Amphiphilic Janus Nanoparticles. ACS Nano 2018, 12 (4), 3646–3657.

18. Poyton, M. F.; Pullanchery, S.; Sun, S.; Yang, T.; Cremer, P. S., Zn(2+) Binds to Phosphatidylserine and Induces Membrane Blebbing. J Am Chem Soc 2020, 142 (43), 18679–18686.

19. Gooran, N.; Tan, S. W.; Frey, S. L.; Jackman, J. A., Unraveling the Biophysical Mechanisms of How Antiviral Detergents Disrupt Supported Lipid Membranes: Toward Replacing Triton X-100. Langmuir 2024, 40 (12), 6524–6536.

20. Kurniawan, J.; Ventrici de Souza, J. F.; Dang, A. T.; Liu, G. Y.; Kuhl, T. L., Preparation and Characterization of Solid-Supported Lipid Bilayers Formed by Langmuir-Blodgett Deposition: A Tutorial. Langmuir 2018, 34 (51), 15622–15639.

21. Crane, J. M.; Kiessling, V.; Tamm, L. K., Measuring lipid asymmetry in planar supported bilayers by fluorescence interference contrast microscopy. Langmuir 2005, 21 (4), 1377–88.

22. Crane, J. M.; Tamm, L. K., Role of cholesterol in the formation and nature of lipid rafts in planar and spherical model membranes. Biophys J 2004, 86 (5), 2965–79.

23. Richter, R. P.; Berat, R.; Brisson, A. R., Formation of solid-supported lipid bilayers: an integrated view. Langmuir 2006, 22 (8), 3497–505.

24. Anderson, T. H.; Min, Y.; Weirich, K. L.; Zeng, H.; Fygenson, D.; Israelachvili, J. N., Formation of supported bilayers on silica substrates. Langmuir 2009, 25 (12), 6997–7005.

25. Groves, J. T.; Ulman, N.; Boxer, S. G., Micropatterning fluid lipid bilayers on solid supports. Science 1997, 275 (5300), 651–3.

26. Reimhult, E.; Hook, F.; Kasemo, B., Vesicle adsorption on SiO2 and TiO2: Dependence on vesicle size. J Chem Phys 2002, 117 (16), 7401–7404.

27. Keller, C. A.; Kasemo, B., Surface specific kinetics of lipid vesicle adsorption measured with a quartz crystal microbalance. Biophys J 1998, 75 (3), 1397–1402.

28. Bruzas, I.; Brinson, B. E.; Gorunmez, Z.; Lum, W.; Ringe, E.; Sagle, L., Surface-Enhanced Raman Spectroscopy of Fluid-Supported Lipid Bilayers. ACS Appl Mater Interfaces 2019, 11 (36), 33442–33451.

29. Mager, M. D.; Almquist, B.; Melosh, N. A., Formation and characterization of fluid lipid bilayers on alumina. Langmuir 2008, 24 (22), 12734–7.

30. Cho, N. J.; Cho, S. J.; Cheong, K. H.; Glenn, J. S.; Frank, C. W., Employing an amphipathic viral peptide to create a lipid bilayer on Au and TiO2. J Am Chem Soc 2007, 129 (33), 10050–1.

31. Naumann, R.; Schiller, S. M.; Giess, F.; Grohe, B.; Hartman, K. B.; Kärcher, I.; Köper, I.; Lübben, J.; Vasilev, K.; Knoll, W., Tethered lipid Bilayers on ultraflat gold surfaces. Langmuir 2003, 19 (13), 5435–5443.

32. Plant, A. L., Supported hybrid bilayer membranes as rugged cell membrane mimics. Langmuir 1999, 15 (15), 5128–5135.

33. Jordan, L. R.; Blauch, M. E.; Baxter, A. M.; Cawley, J. L.; Wittenberg, N. J., Influence of brain gangliosides on the formation and properties of supported lipid bilayers. Colloids Surf B Biointerfaces 2019, 183, 110442.

34. Sendecki, A. M.; Poyton, M. F.; Baxter, A. J.; Yang, T.; Cremer, P. S., Supported Lipid Bilayers with Phosphatidylethanolamine as the Major Component. Langmuir 2017, 33 (46), 13423–13429.

35. Sundh, M.; Svedhem, S.; Sutherland, D. S., Influence of phase separating lipids on supported lipid bilayer formation at SiO2 surfaces. Phys Chem Chem Phys 2010, 12 (2), 453–60.

36. Tabaei, S. R.; Choi, J. H.; Haw Zan, G.; Zhdanov, V. P.; Cho, N. J., Solvent-assisted lipid bilayer formation on silicon dioxide and gold. Langmuir 2014, 30 (34), 10363–73.

37. Hohner, A. O.; David, M. P.; Radler, J. O., Controlled solvent-exchange deposition of phospholipid membranes onto solid surfaces. Biointerphases 2010, 5 (1), 1–8.

38. Ferhan, A. R.; Yoon, B. K.; Park, S.; Sut, T. N.; Chin, H.; Park, J. H.; Jackman, J. A.; Cho, N. J., Solvent-assisted preparation of supported lipid bilayers. Nat Protoc 2019, 14 (7), 2091–2118.

39. Tabaei, S. R.; Jackman, J. A.; Kim, S. O.; Liedberg, B.; Knoll, W.; Parikh, A. N.; Cho, N. J., Formation of cholesterol-rich supported membranes using solvent-assisted lipid self-assembly. Langmuir 2014, 30 (44), 13345–52.

40. Tae, H.; Yang, C.; Cho, N. J., Artificial Cell Membrane Platforms by Solvent-Assisted Lipid Bilayer (SALB) Formation. Accounts Mater Res 2022, 3 (12), 1272–1284.

41. Su, H.; Liu, H. Y.; Pappa, A. M.; Hidalgo, T. C.; Cavassin, P.; Inal, S.; Owens, R. M.; Daniel, S., Facile Generation of Biomimetic-Supported Lipid Bilayers on Conducting Polymer Surfaces for Membrane Biosensing. ACS Appl Mater Interfaces 2019, 11 (47), 43799–43810.

42. Timson, D. J., The roles and applications of chaotropes and kosmotropes in industrial fermentation processes. World J Microbiol Biotechnol 2020, 36 (6), 89.

43. Sachs, J. N.; Woolf, T. B., Understanding the Hofmeister effect in interactions between chaotropic anions and lipid bilayers: molecular dynamics simulations. J Am Chem Soc 2003, 125 (29), 8742–3.

44. Cray, J. A.; Russell, J. T.; Timson, D. J.; Singhal, R. S.; Hallsworth, J. E., A universal measure of chaotropicity and kosmotropicity. Environ Microbiol 2013, 15 (1), 287–96.

45. Senthilkumar, R.; Sharma, K. K., Effect of chaotropic agents on the structure-function of recombinant acylpeptide hydrolase. J Protein Chem 2002, 21 (5), 323–32.

46. Julien, J. A.; Rousseau, A.; Perone, T. V.; LaGatta, D. M.; Hong, C.; Root, K. T.; Park, S.; Fuanta, R.; Im, W.; Glover, K. J., One-step site-specific S-alkylation of full-length caveolin-1: Lipidation modulates the topology of its C-terminal domain. Protein Sci 2023, 32 (11), e4791.

47. Oku, N.; MacDonald, R. C., Solubilization of phospholipids by chaotropic ion solutions. J Biol Chem 1983, 258 (14), 8733–8.

48. Oku, N.; Macdonald, R. C., Formation of Giant Liposomes from Lipids in Chaotropic Ion Solutions. Biochimica Et Biophysica Acta 1983, 734 (1), 54–61.

49. Jonsson, P.; Jonsson, M. P.; Tegenfeldt, J. O.; Hook, F., A method improving the accuracy of fluorescence recovery after photobleaching analysis. Biophys J 2008, 95 (11), 5334–48.

50. Cho, N. J.; Frank, C. W.; Kasemo, B.; Hook, F., Quartz crystal microbalance with dissipation monitoring of supported lipid bilayers on various substrates. Nat Protoc 2010, 5 (6), 1096–106.

51. Tabaei, S. R.; Jackman, J. A.; Kim, S. O.; Zhdanov, V. P.; Cho, N. J., Solvent-assisted lipid self-assembly at hydrophilic surfaces: factors influencing the formation of supported membranes. Langmuir 2015, 31 (10), 3125–34.

52. Dixon, M. C., Quartz crystal microbalance with dissipation monitoring: enabling real-time characterization of biological materials and their interactions. J Biomol Tech 2008, 19 (3), 151–8.

53. Jackman, J. A.; Tabaei, S. R.; Zhao, Z.; Yorulmaz, S.; Cho, N. J., Self-assembly formation of lipid bilayer coatings on bare aluminum oxide: overcoming the force of interfacial water. ACS Appl Mater Interfaces 2015, 7 (1), 959–68.

54. Gunko, V. M.; Turov, V. V.; Zarko, V. I.; Voronin, E. F.; Tischenko, V. A.; Dudnik, V. V.; Pakhlov, E. M.; Chuiko, A. A., Active site nature of pyrogenic alumina/silica and water bound to surfaces. Langmuir 1997, 13 (6), 1529–1544.

55. Turov, V. V.; Leboda, R., Application of 1H NMR spectroscopy method for determination of characteristics of thin layers of water adsorbed on the surface of dispersed and porous adsorbents. Adv Colloid Interface Sci 1999, 79 (2-3), 173–211.

56. Zwang, T. J.; Fletcher, W. R.; Lane, T. J.; Johal, M. S., Quantification of the layer of hydration of a supported lipid bilayer. Langmuir 2010, 26 (7), 4598–601.

