## Supporting Information for "Chaotropic Agent-assisted Supported Lipid Bilayer Formation"

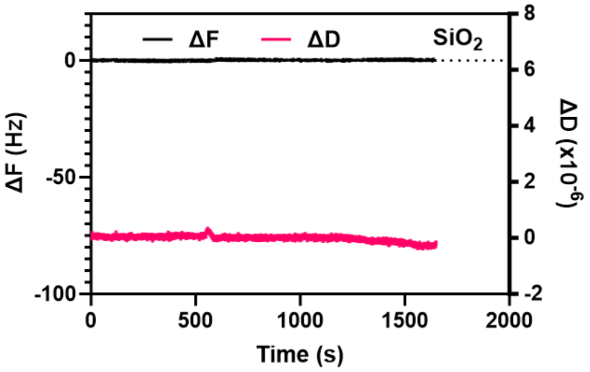


**Figure S1.** Negligible BSA adsorption on POPC SLB formed by vesicle rupture. The frequency (ΔF) and dissipation (ΔD) shifts are shown in black and magenta, respectively.
